# Effects of transcranial magnetic stimulation on the human brain recorded with intracranial electrocorticography: First-in-human study

**DOI:** 10.1101/2022.01.18.476811

**Authors:** Jeffrey B. Wang, Joel E. Bruss, Hiroyuki Oya, Brandt D. Uitermarkt, Nicholas T. Trapp, Phillip E. Gander, Matthew A. Howard, Corey J. Keller, Aaron D. Boes

## Abstract

Transcranial magnetic stimulation (TMS) is increasingly used as a noninvasive technique for neuromodulation in research and clinical applications, yet its mechanisms are not well understood. Here, we present the first in-human study evaluating the effects of TMS using intracranial electrocorticography (iEEG) in neurosurgical patients. We first evaluated safety in a gel-based phantom. We then performed TMS-iEEG in 20 neurosurgical participants with no adverse events. Next, we evaluated brain-wide intracranial responses to single pulses of TMS to the dorsolateral prefrontal cortex (dlPFC) (N=10, 1414 electrodes). We demonstrate that TMS preferentially induces neuronal responses locally within the dlPFC at sites with higher electric field strength. Evoked responses were also noted downstream in the anterior cingulate and anterior insular cortex, regions functionally connected to the dlPFC. These findings support the safety and promise of TMS-iEEG in humans to examine local and network-level effects of TMS with higher spatiotemporal resolution than currently available methods.

## Introduction

Transcranial magnetic stimulation (TMS) is a noninvasive technique for modulating the regional excitability of the human brain (Dayan et al., 2016; Hallett, 2007). Clinically, it is FDA cleared for depression, smoking cessation, migraines, and obsessive compulsive disorder, with clinical trials underway for many other neuropsychiatric disorders (Elias et al., 2021; Lefaucheur et al., 2020). It is also increasingly used as a neuroscientific experimental tool to probe neural circuitry within the human brain. The neurophysiological effects of TMS in animal models has been investigated extensively, demonstrating that TMS induces local neuronal firing within milliseconds of the delivered TMS pulse (Romero et al., 2019). However, efforts to understand the physiological effects of TMS in humans have been hampered by methodological limitations (Boes et al., 2018a; Chervyakov et al., 2015), specifically the lack of either spatial or temporal resolution, as is the case with surface EEG and fMRI, respectively. As such, there is a critical need for novel methods that can map the effects of TMS with high spatial and temporal resolution simultaneously. This would facilitate further insights into the neural mechanisms of TMS, guiding optimization of its use as a research tool and treatment for neuropsychiatric disorders. Specifically, elucidating the underlying neural mechanisms can further facilitate treatment improvement for major depression and other psychiatric disorders above the current moderate (∼30-50%) clinical response.

Neural activity can be measured with high spatiotemporal resolution from intracranial electroencephalography (iEEG) recorded from neurosurgical epilepsy patients using electrodes either implanted within the brain or on its surface. iEEG has been used to delineate the temporal dynamics and spatial spread following intracranial electrical stimulation (Huang et al., 2019; Keller et al., 2014a, 2018; Matsumoto et al., 2004, 2012) and is a promising tool for providing similar resolution following non-invasive neuromodulatory techniques. Indeed, recent work with non-invasive transcranial direct and alternating current stimulation (tDCS & tACS) in humans have shown the utility of investigating these effects with iEEG (Chhatbar et al., 2018; Huang et al., 2019; Lafon et al., 2017; Opitz et al., 2016; Vöröslakos et al., 2018). These studies demonstrated that higher stimulation amplitude than is typically used may be needed to reliably induce intracranial effects (Vöröslakos et al., 2018). Moreover, protocols that were presumed to drive specific oscillation frequencies did not find supporting evidence from iEEG (Lafon et al., 2017). To date these iEEG studies have not been extended to TMS, though studies of TMS with intracranial recordings have been highly informative in nonhuman primates (Mueller et al., 2014; Romero et al., 2019). If applied to humans, these data acquired with high spatiotemporal resolution could help to better characterize the effects of TMS on the human brain, illuminating both local responses induced directly by TMS and downstream network-level responses propagated to connected brain regions.

Current consensus guidelines on TMS safety include intracranial hardware as a contraindication for TMS administration (Rossi et al., 2009; Rossini et al., 2015). Emerging safety data relevant to using TMS in the setting of intracranial hardware has been encouraging (Gaynor et al., 2008; Kühn et al., 2002; Kumar and Chen, 1999; Phielipp et al., 2017; Udupa et al., 2016); see Rossi et al., (2021) for a more extensive review. Recent animal studies have demonstrated that TMS can be applied safely in the presence of iEEG (Romero et al., 2019), but the safety of this technique has not been evaluated in humans. To date there has not been an extensive study of the safety of TMS applied directly to intracranial electrodes, nor has there been an investigation of the electrophysiological effects of TMS using iEEG.

In this study we first investigate the safety of applying TMS with iEEG by conducting experiments in a gel-based phantom brain. We evaluate whether heating, displacement, or the induction of secondary currents in the intracranial electrodes would pose safety risks that would preclude further investigation of this approach in humans. After demonstrating safety using the gel-based phantom, we show that TMS has a favorable safety profile based on in-vivo combined TMS and iEEG (TMS-iEEG) in 20 patients. Next, in a subset of 10 patients, we evaluate the local and downstream electrophysiological effects of single pulses of TMS applied to the dorsolateral prefrontal cortex (dlPFC), the therapeutic target for depression and other neuropsychiatric disorders (Perera et al., 2017). Here, we demonstrate that these single TMS pulses induce neuronal responses both locally within the dlPFC and in regions functionally connected to the stimulation site, including deep structures such as the anterior cingulate cortex (ACC). Together, these findings show that 1) TMS-iEEG is a viable tool for studying the electrophysiological effects of TMS on the human brain and that 2) TMS induces responses both locally and in functionally connected downstream neuronal populations in humans.

## Online content

### Methods

#### Safety testing using a gel-based phantom brain

We addressed the safety of performing TMS with iEEG by first using a gel-based phantom brain as a model. The analyses were focused on evaluating three main concerns: 1) the possibility of electrodes heating, 2) electrode displacement, which could damage surrounding tissue, and 3) induction of secondary electric currents in the intracranial electrodes from the time-varying magnetic field generated by TMS. To evaluate these possibilities, we delivered TMS to a gel phantom with intracranial electrodes placed within the gel and on the surface to mimic human experimental conditions, as used previously to investigate safety related to experiments with intracranial electrodes (Oya et al., 2017). The distance from the TMS coil to the electrode contacts was set to 10 mm, which conservatively approximates the smallest distance possible (and thus the highest amplitude in magnetic field) between the coil and iEEG electrodes in human experiments, where TMS must cross the skin, skull, and cerebrospinal fluid space prior to reaching the electrodes (Davis, 2021; Lu and Ueno, 2017).

##### TMS Equipment & Stimulation Parameters

TMS equipment included a MagVenture MagVita X100 230V system with a figure-of-eight liquid-cooled Cool-B65 A/P coil (MagVenture; Alpharetta, GA, USA). Stimulation pulse was biphasic sinusoidal with a pulse width of 290 microseconds for all experiments except for those measuring displacement, which was monophasic to avoid the possible cancellation of displacing forces. For safety testing in the phantom brain, the stimulation intensity range was set to 100% machine output delivered at 10 – 40 Hz. For in vivo experiments, the stimulator output was delivered at a percentage of each subject’s individually tested motor threshold (see Table 1) at a frequency of 0.5 Hz for the ‘single pulse’ dlPFC analyses. Other stimulation parameters in the assessment of safety included 10 Hz, 20 Hz and intermittent theta burst stimulation (iTBS) patterns.

**Table 1.** TMS stimulation parameters, electrode characteristics, and imaging information.

| ID | Stimulation Site | MT (% machine output) | Stimulation Intensity (% MT) | Resting State (min) | Implantation Type | # Electrodes (# Removed) | Day of Testing Medications |
| --- | --- | --- | --- | --- | --- | --- | --- |
| 429 | Left & Right dlPFC | 86% | 100% | 0 | ECoG | 199 (27) | albuterol sulfate, folic acid, lorazepam, norgestimate-ethinyl estradiol, sertraline HCl |
| 430 | Left dlPFC | 53% | 100% | 14.3 | sEEG | 72 (30) | clonazepam, lamotrigine, risperidone, topiramate, venlafaxine HCl |
| 460 | Left dlPFC | 48% | 120% | 14.5 | ECoG | 185 (13) | aspirin, cetirizine HCl, doxazosin mesylate, escitalopram oxalate, lamotrigine, losartan potassium, magnesium chloride, riboflavin, tamsulosin HCl |
| 477 | Left Parietal; Left dlPFC | 59% | 120% | 23.8 | ECoG | 148 (13) | albuterol sulfate, clonazepam, fluticasone propion/salmeterol, ipratropium/albuterol sulfate, levetiracetam, montelukast sodium, oxcarbazepine, sertraline HCl, topiramate |
| 483 | Left Parietal; Left dlPFC | 75% | 120% | 17.2 | sEEG | 226 (57) | cefazolin, clobazam, felbamate, insulin, miralax |
| 518 | Left dlPFC | ** | ** | 23.8 | sEEG | 216 (62) | clonazepam, lamotrigine, topiramate |
| 534 | Left dlPFC | * | 40%, 50%, & 70% machine output*** | 0 | sEEG | 224 (7) | clobazam, oxcarbazepine |
| 538 | Left dlPFC | 80% | 80% & 100% | 24.5 | sEEG | 198 (18) | buspirone, cetirizine, diphenhydramine, docusate, famotidine, levetiracetam, lorazepam, vancomyzine |
| 559 | Left dlPFC | 73% | 80%, 100%, & 110% | 24.0 | sEEG | 145 (18) | carbamazepine, clonazepam, lacosomide, levetiracetam, ondansetron, sumatriptan, zonisamide |
| 561 | Left dlPFC | * | 40%, 50%, & 70% machine output*** | 24 | sEEG | 242 (28) | acetaminophen, carbamazepine, cefazolin, cetirizine, docusate, ibuprofen, lacosamide, lorazepam, ondansetron, oxycodone, miralax, sertraline |

Abbreviations: dlPFC Dorsolateral Prefrontal Cortex, MT Motor Threshold, HCl Hydrogen Chloride ECoG electrocorticography, sEEG stereoelectroencephalography \*not formally evaluated; \*\* motor cortex inaccessible; \*\*\*reporting percent machine output as motor cortex inaccessible <sup>+</sup> Subject 483 had a multiband resting state acquisition: TR = 1000 ms, TE = 30 ms, Flip Angle = 65 deg. Voxel size = 2.44 x 2.44 x 2.5 mm, 4 blocks of 10-minute gradient echo EPI (1032 volumes). Subjects 429 and 561 were excluded from dlPFC analyses due to amplifier saturation in >10% of electrodes.

##### Gel Phantom Apparatus and Electrodes

We used a custom-made gel phantom filled with polyacrylic acid saline gel placed in an 8-inch cubic container with a 3/16-inch polymethyl methacrylate wall. It has similar physical, thermal and electrical properties to the human brain. Its fabrication details are described by the American Society for Testing and Materials standards section F2182 (Committee F04 on Medical and Surgical Materials and Devices, 2011). The electrodes included a 32-contact grid electrode and an 8-contact penetrating depth electrode array with 1-cm spacing, made of non-ferromagnetic platinum (SD08R-SP10X-000 and a 4 connector L-SRL-8DIN; Ad-Tech; Racine, WI, USA), all of which are embedded in a silicon-based sheet (silastic) material. Inter-contact impedance within the gel phantom at 100 Hz was 2.82 +/- 1.1 kiloohms (mean and standard deviation) and at 1000 Hz was 1.44 +/- 0.87 kiloohms.

##### Temperature Measurements

Fiberoptic fluorescent temperature sensors were used to measure local temperature changes that may occur in gel-implanted electrodes (FTX-300 optic signal conditioner and PRBMR1 optical fiber probes; OSENSA innovations; BC, Canada). Temperature measurements were obtained from 2 sensors simultaneously located at 10 and 20 mm from the coil at a 2 Hz sampling rate with accompanying time stamps. The temperature sensing probes were mechanically attached perpendicular to the metallic electrodes to maintain direct physical contact. The sensor cables were secured to the electrode by using medical tape and collodion. TMS was delivered at 100% machine output at 10, 20 and 40 Hz while temperature was recorded continuously for 350 seconds (5.8 minutes).

##### Video Monitoring of Electrode Displacement

Current flow across electrodes induced by TMS could produce a secondary magnetic field that would interact with the primary magnetic field produced by the TMS coil, potentially displacing the electrode contacts. To assess this effect, we used a Sony HDR-XR200 to record subdural grid and depth electrodes suspended in normal saline within a beaker at a frame rate of 30 frames-per-second. Monophasic TMS was administered to the electrodes at 40 Hz, 100% machine output at less than 10 mm. Video images of the electrodes were recorded with a resolution that allows visualization of electrode displacement as small as 1 mm.

##### Induced Current Measurement

We measured the induced voltages across pairs of electrode contacts. The voltage measurements were obtained using an oscilloscope probe (10x attenuation passive voltage probe) with peak-to-peak voltage measured (Tektronix TDS2022 200 MHz scope; Beaverton, OR, USA and Agilent 54642D 500 MHz; Santa Clara; CA; USA). Parallel and perpendicular orientations of the depth electrodes were tested along with various positions across the surface of the TMS coil. Induced voltage in cable and connectors was also tested.

#### TMS in neurosurgical patients

##### Subjects

20 neurosurgical patients with medically intractable epilepsy participated in this study and were included in the safety analysis (10 Females, age range 13 - 56, mean 28 +/- 13). A subset of 10 patients (5 Females, age range 14 – 52 years, mean 25 +/-11 SD) received ‘single pulse’ TMS to the dlPFC (pulses applied at 0.5 Hz) and were selected for the main analysis quantifying the evoked response after dlPFC TMS. Each patient was admitted to the University of Iowa Hospitals and Clinics for 14 days of monitoring with intracranial electrodes to localize their seizure focus (Table 2, Supplementary Table 1,2). An iEEG monitoring plan was generated by clinicians from the University of Iowa Comprehensive Epilepsy Program. All electrodes were placed solely on the basis of clinical requirements to identify seizure foci (Nagahama et al., 2018). Following intracranial electrode implantation surgery patients remained in an electrically shielded epilepsy monitoring room in the University of Iowa’s Clinical Research Unit. TMS experiments were conducted after the final surgical treatment plan was agreed upon between the clinical team and the patient, typically 1-2 days before the planned electrode explantation operation and 24 hours after the patient had restarted anti-epileptic medications. All experimental procedures were approved by the University of Iowa Institutional Review Board, who had available our gel phantom safety experiments prior to reviewing the IRB for human subjects. Written informed consent was obtained from all participants.

**Table 2.**
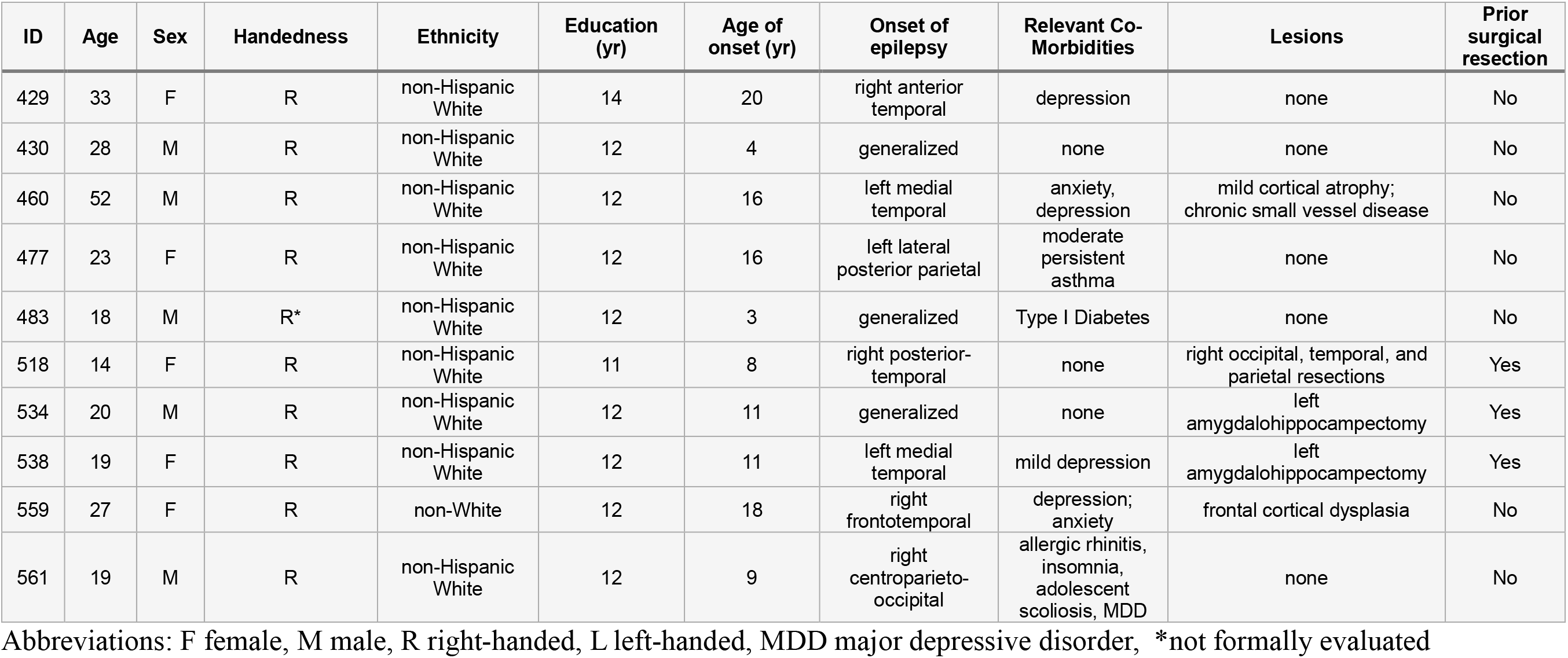
Patient Demographics

| ID | Age | Sex | Handedness | Ethnicity | Education (yr) | Age of onset (yr) | Onset of epilepsy | Relevant Co-Morbidities | Lesions | Prior surgical resection |
| --- | --- | --- | --- | --- | --- | --- | --- | --- | --- | --- |
| 429 | 33 | F | R | non-Hispanic White | 14 | 20 | right anterior temporal | depression | none | No |
| 430 | 28 | M | R | non-Hispanic White | 12 | 4 | generalized | none | none | No |
| 460 | 52 | M | R | non-Hispanic White | 12 | 16 | left medial temporal | anxiety, depression | mild cortical atrophy; chronic small vessel disease | No |
| 477 | 23 | F | R | non-Hispanic White | 12 | 16 | left lateral posterior parietal | moderate persistent asthma | none | No |
| 483 | 18 | M | R* | non-Hispanic White | 12 | 3 | generalized | Type I Diabetes | none | No |
| 518 | 14 | F | R | non-Hispanic White | 11 | 8 | right posterior-temporal | none | right occipital, temporal, and parietal resections | Yes |
| 534 | 20 | M | R | non-Hispanic White | 12 | 11 | generalized | none | left amygdalohippocampectomy | Yes |
| 538 | 19 | F | R | non-Hispanic White | 12 | 11 | left medial temporal | mild depression | left amygdalohippocampectomy | Yes |
| 559 | 27 | F | R | non-White | 12 | 18 | right frontotemporal | depression; anxiety | frontal cortical dysplasia | No |
| 561 | 19 | M | R | non-Hispanic White | 12 | 9 | right centroparieto-occipital | allergic rhinitis, insomnia, adolescent scoliosis, MDD | none | No |

Abbreviations: F female, M male, R right-handed, L left-handed, MDD major depressive disorder, \*not formally evaluated

##### Pre-Implantation Neuroimaging

Prior to implantation of intracranial electrodes, patients underwent an anatomical and functional MRI scan within two weeks of the electrode implantation surgery. The scanner was a 3 Tesla GE Discovery MR750W with a 32-channel head coil (GE; Boston, MA, USA). The pre-electrode implantation anatomical T1 scan was obtained with following parameters: 3d FSPGR BRAVO sequence: FOV = 25.6 cm, Flip angle = 12 deg., TR = 8.50 ms, TE = 3.288 ms, Inversion time = 450 ms, voxel size = 1.0 x 1.0 x 0.8 mm. Resting-state functional MRI (rs-fcMRI) was also obtained. Participants were asked to keep their eyes open and a fixation cross was presented through a projector. Five blocks of 5-minute gradient-echo EPI runs were acquired (650 volumes) with the following parameters: FOV = 22.0 cm, TR = 2260 ms, TE = 30 ms, Flip Angle = 80 deg., voxel size = 3.45 x 3.45 x 4.0 mm. In some cases, fewer resting state acquisition sequences were used in the final analysis due to movement artifact or the full scanning session was not completed (Table 1). For each patient, rs-fcMRI runs were acquired in the same session but non-contiguously (dispersed within an imaging session to avoid habituation).

##### Post-Implantation Imaging

The day following the electrode implantation surgery, subjects underwent a second MRI and thin-slice volumetric computerized tomography (CT) scans obtained for clinical purposes. The MRI was performed on a Siemens Skyra 3T scanner acquiring an MPRAGE sequence with the following parameters: FOV = 25.6 cm, TR = 1900 ms, TE = 3.44 ms, TI = 1000 ms, Flip Angle = 10 deg., Voxel size = 1.0 x 1.0 x 1.0 mm. The CT scan was obtained with a voxel size of 0.47 x 0.47 x 1.0 mm. These post-implantation scans were used to identify the position of each electrode contact, as described next.

##### Image Processing and Intracranial Electrode Localization

For each subject, T1 and T2 MRI sequences were co-registered via an affine transform using FSL’s FLIRT software (Jenkinson and Smith, 2001; Jenkinson et al., 2002). The location of each electrode was identified on the post-implantation MRI T1-weighted image and CT, aided by cross-referencing intraoperative photographs when possible (surface electrodes). Once electrode contact locations were identified as accurately as possible, the post-implantation scans were transformed to the pre-implantation T1 anatomical space with nonlinear three-dimensional thin-plate spline warping. This process was aided by 50-100 manually placed control points that help to accurately align scans while accounting for the possibility of post-surgical brain shift (Oya et al., 2017, 2018). Reconstruction of the anatomical locations of the implanted electrodes and their mapping onto a standardized set of coordinates across subjects was performed using FreeSurfer image analysis suite (Version 6.0; Martinos Center for Biomedical Imaging, Charleston, MA; Dale et al., 1999). Each FreeSurfer run was reviewed for anatomical accuracy and corrected or re-ran as needed. Coordinates were obtained for each contact in native space on the pre-implantation MRI, pial surface, along with CIT168 and MNI152 space, and the transform between these spaces was calculated using ANTs’ antsRegistration (Klein et al., 2009), and FSL’s FNIRT with FEAT processing. FieldMap data was incorporated with BBR registration to calculate an EPI to T1 nonlinear transformation. Warp fields were then summed, and inverse transforms were created to facilitate bidirectional EPI to MNI152 transforms. The Desikan-Killiany-Tourville (DKT) atlas within the FreeSurfer package was used as an anatomical reference for electrode locations (Desikan et al., 2006).

##### Resting State Functional Connectivity MRI

Standard preprocessing was applied to the rs-fcMRI data acquired in the pre-implantation scan using FSL’s FEAT pipeline, including de-spiking, slice timing correction, and spatial alignment. White matter and ventricles were masked using a set of ROIs generated in MNI152 space and transformed to each subject’s EPI scan. Nuisance regressors, included: 1) Global signal, 2) White matter, 3) CSF, and 4) 6 motion parameters (3 rotations and 3 translations). They were extracted from these masks and detrended with second order polynomials. Temporal bandpass filtering was 0.008–0.08 Hz. Regression was performed using the time series calculated above as well as motion parameters and their derivatives. Spatial smoothing was applied with a 6 mm full-width at half maximum Gaussian kernel. The first 2 images from each run were discarded. Frame censoring was applied when the Euclidean norm of derivatives of motion parameters exceeded 0.3 mm (Power et al., 2012). All runs were processed in native EPI space and then concatenated in MNI152 space.

Seed-based rs-fcMRI analysis was performed to evaluate functional connectivity between the stimulation site for TMS and the rest of the brain. The stimulation site was estimated at the point where a vector extending from the coil intersected with the nearest vertex on the cerebral cortex. The vector was calculated extending from the center of the figure-of-eight coil based on its three-dimensional position during stimulation. Directional cosines derived from neuronavigation Brainsight software (Rogue Research, Quebec, CA) were used to create this vector which was then intersected with a FreeSurfer-derived mesh of the scalp to find the closest surface coordinate to the center of the coil. Pial coordinates of the stimulation site were converted from subject vertex to MNI152 vertex, then RAS coordinates and 4mm diameter spherical ROIs were generated using AFNI’s 3dUndump (Cox, 1996). The mean blood oxygen level dependent (BOLD) signal time series was extracted from each stimulation site ROI and from all other voxels throughout the brain. Note that the rs-fcMRI data was acquired in the pre-implantation scan and the post-implantation scans were used to identify the electrode contact locations, with co-registration of the post-implantation to pre-implantation scan using ANTs. Seed-based functional connectivity maps were created by calculating Pearson’s correlation coefficients between the time series of the stimulation site and all other voxels in the brain, with a Fisher r-to-Z transform. Group resting state functional connectivity MRI (rs-fcMRI) maps were also generated from each stimulation site (transforming to MNI152 space) using publicly available normative rs-fcMRI data from 98 healthy individuals, using the same processing as was described previously (Boes et al., 2018b; Holmes et al., 2015).

##### Transcranial Magnetic Stimulation

The TMS experiment was conducted 12-13 days post-implantation surgery and after restarting anti-seizure medicine. TMS was performed with a Cool-B65 Active/Placebo (A/P) liquid cooled butterfly coil using the same MagVenture system described above. Neuronavigation using frameless stereotaxy was guided with Brainsight software supplied with the pre-implantation T1 / MPRAGE anatomical scan. Stimulation parameters of each TMS pulse (location, coil position, and dI/dt) were recorded in Brainsight during all experimental trials. Motor threshold procedures were performed for each participant prior to experimental testing. The hand knob of the motor cortex was identified from the MRI and used as a starting target for motor threshold testing. The starting intensity was 30% machine output and adjusted in 5-10% increments until hand movements were observed in 50% of trials.

The main analysis included single pulses of TMS delivered at 0.5 Hz to the dlPFC to evaluate the strength and distribution of evoked responses. Pulses were delivered at 0.5 Hz at 100% or 120% of motor threshold. 100% motor threshold was utilized if 120% was not tolerated by the participant due to pain. The anatomical target of stimulation was the dorsolateral prefrontal cortex (dlPFC) defined by the Beam F3 region (Beam et al., 2009), identified by transforming published coordinates (MNI 1mm: -41.5, 41.1, 33.4 (Fried et al., 2014)) into each subject’s native T1 and displaying it in Brainsight. The stimulation site was modified slightly in some cases if access was impeded by head wrap or anchor bolts for securing electrodes. The motor threshold and locations of TMS delivery are provided in Table 1. To evaluate site specificity of TMS evoked potentials, we applied the identical simulation protocol of 0.5 Hz single pulses of TMS to the inferior parietal lobe at a site defined by functional connectivity to the hippocampus (Wang et al., 2015) in a subset of participants (N=2). To evaluate neural responses that may be due to auditory effects (Poorganji et al., 2021), we applied sham TMS to the dlPFC, with the TMS coil (Cool-B65 A/P) flipped 180-degrees such that the magnetic field was directed away from the head. Additional details and any subject-specific deviations from these parameters are described in Supplementary Tables 1 and 2. Other participants that did not receive 0.5 Hz dlPFC stimulation were included in the assessment of TMS-iEEG safety. These subjects received alternate stimulation protocols, with details provided in Supplementary Table 2.

##### iEEG Recording

Electrode implantation and recording protocols have previously been described in detail (Gander et al., 2019; Nourski and Howard, 2015). Depth and grid electrode arrays were manufactured by Ad-Tech Medical (Racine, WI, USA). Depth arrays (platinum macro-contacts, with 5- or 10-mm inter-contact spacing) were stereotactically implanted for each subject; grid arrays (platinum-iridium, 2.3 mm exposed diameter, with 5- or 10-mm inter-contact spacing) were placed on the cortical surface. A platinum-iridium strip electrode placed in the midline subgaleal space was used as a reference. iEEG data acquisition was controlled by a TDT RZ2 real-time processor (Tucker-Davis Technologies; Alachua, FL, USA). Collected iEEG data were amplified, filtered (ATLAS, Neuralynx, Bozeman, MT; 0.7-800 Hz bandpass, 12 dB/octave rolloff), digitized at a sampling rate of 8000 Hz, and stored for subsequent offline analysis. In all subjects, contacts were excluded from analysis if they were determined to be involved in either the generation or early propagation of seizures, were implanted outside the brain or within white matter, or if artifact saturated the amplifier. Subjects were removed from analysis if TMS artifact saturated the amplifier in >10% of contacts, which occurred in 2 out of 10 subjects.

##### Intracranial Stimulation

To evaluate if evoked responses were sensory in nature, we compared responses from TMS to those elicited from single pulse direct electrical stimulation delivered from intracranial electrodes on the surface of the dlPFC, as this method produces minimal perceptual effects. Intracranial direct electrical stimulation has been described previously from our groups (Keller et al., 2011, 2014b, 2014a; Rocchi et al., 2021). Briefly, 50 single constant current electrical stimulation pulses (biphasic charge-balanced square wave, duration = 0.2 ms/phase, 9 or 12 mA; 2s inter-stimulus interval) were applied in bipolar configuration through AM Systems Model 2200 (Sequim, WA, USA) stimulus isolators connected to adjacent intracranial electrode pairs (Ad-Tech Medical Instrument Corp.). Other aspects of intracranial recordings were identical to those described above (*iEEG recording*). The coordinates of the stimulated dlPFC contacts for each subject can be found in Fig. 4 legend. No subjects showed evidence of or voiced any pain or discomfort during intracranial electrical stimulation.

#### Electrophysiology analysis

##### Preprocessing of iEEG Data

Data preprocessing and analysis was performed offline using the FieldTrip toolbox (Oostenveld et al., 2011) and with custom scripts (MATLAB; Mathworks; Portola Valley, CA, USA). Artifact rejection consisted of three stages: line noise removal, TMS stimulation pulse artifact removal, and subsequent decay artifact removal, as standard in previous TMS preprocessing algorithms (Rogasch et al., 2017; Wu et al., 2018). First, line noise (60 Hz) and up to seven harmonics were removed using a notch filter (3^rd^ order Butterworth filter, cutoff frequencies 57-63 Hz). Seven harmonics were chosen based on visual inspection of the spectrogram prior to artifact removal. Removal of frequencies at this high range were necessary because the TMS artifact consists of frequencies above 300 Hz and at times overlapped with higher harmonics of line noise, so a low-pass filter is insufficient during this stage. Next, TMS-induced stereotyped stimulation pulse artifacts ∼15ms in duration that reached input range saturation were removed by replacing the stimulation artifact with stationary iEEG time series that represented similar amplitude and spectral profile as the background signal. This procedure has been detailed previously (Crowther et al., 2019) and is preferred over simple spline interpolation given short intervals between pulses to circumvent the possibility of introducing large spectral changes. For each channel, we extracted iEEG signal with equal length as the stimulation artifact immediately preceding and following the artifact. We reversed the iEEG signal and applied a tapering matrix (1:1/n:0 for the preceding data, 0:1/n:1 for the following data, where n is the number of samples contained in the artifact). The two iEEG signals were added together and subsequently used to replace the artifact period.

After stimulation pulse artifact removal, the longer decay artifact occurring over ∼150 ms was removed using the Adaptive Detrend Algorithm (ADA) as previously described (Casula et al., 2017). First, each stimulation artifact-scrubbed epoch was restricted from 15-500 ms. Each subsequent epoch was then fit with both a double exponential and linear model, defined as

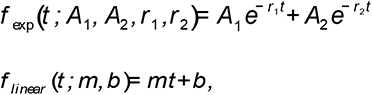

where *t* is the time in ms and *A*_1_, *A*_2_, *r*_1_, *r*_2_, *m*, *b* are fitted parameters. After fitting, performance for both models were quantified using the Akaike Information Criterion (AIC), defined as

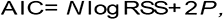

where *N* is the number of data points, *RSS* is the sum of the squares of the residuals, and *P* is the number of fitted parameters in the model. The model with the lowest AIC was chosen and subtracted from the corresponding signal. Combined application of these two models is necessary to address both linear and exponential decay artifacts, which have been previously reported in the literature (Casula et al., 2017).

After artifact rejection, data was then filtered from 1-35 Hz (second order Butterworth filter) to isolate the slower evoked potential within the delta to low gamma bands. Finally, data was downsampled to 300Hz, epoched from -1000ms to 500ms, and baseline corrected to the pre-stimulus voltage between -250ms to -50ms. As a negative control, random 1500 ms epochs were created from the baseline (i.e. prior to TMS stimulation) data by choosing random start times during the baseline period from a uniform distribution. iEEG data during sham TMS was processed in an identical manner as active TMS to act as an additional negative control.

##### Significance Testing and Quantification of iTEPs

To examine and quantify evoked potentials after TMS delivery, we utilized nonparametric clustering as previously described (Maris and Oostenveld, 2007). We calculated one-sample *t* statistics at every time point from 15-100 ms after stimulation to form clusters of significant time points based on temporal adjacency at an alpha level of 0.05 (Maris and Oostenveld, 2007). A cluster was defined as a contiguous region of significant time points with the same t-score polarity. A cluster’s “total statistic” was calculated as the sum of the t-statistic within the cluster. Our final cluster-level statistic was obtained by taking the sum of the three largest individual cluster statistics. To generate the null distribution, we calculated the cluster *t* statistic for randomly shuffled iEEG signals based on 1000 simulations. The true cluster *t* statistic was compared with this null distribution and the evoked potential was considered significant using a *p* value of 0.05 after FDR correction (Yekutieli and Benjamini, 1999) for multiple contacts comparison.

With this method, we performed three pairs of comparison: *TMS vs*. *baseline*, *sham vs*. *baseline*, and *TMS vs*. *sham*. As the active TMS compared to sham condition was of interest, we defined a channel to have a significant TMS-specific neural response if it met the following criteria: 1) TMS was significantly different from both the baseline and sham conditions; 2) TMS response exceeded 10 μV; and 3) Sham was not significantly different from baseline. Sham vs. baseline was required in significance testing as there were contacts where sham and TMS both elicited significant responses but at different amplitudes. Requiring a TMS-evoked potential to be significantly different from both baseline and sham lowered the relative probability of single deflections due to artifact or noise from falsely being classified as a significant evoked potential. The specific threshold of 10 μV for TMS response was chosen following visual inspection of noise across participants, which was consistent from subject to subject.

##### Visualizing the Spatial Distribution of Significant iTEPs

To visualize the location of TMS-specific evoked responses across subjects we generated color-coded ‘heat’ maps depicting the percentage of electrodes in the region showing TMS-specific evoked responses. Specifically, we imported the MNI152 pial surface from FreeSurfer and for each vertex *i* with coordinate *v_i_* we defined the percentage index

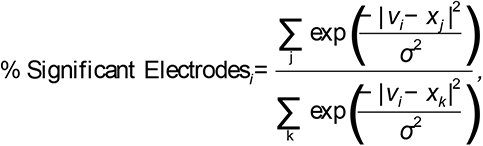

where *j* is the set of significant electrodes, *k* is the set of all electrodes, *x* is the MNI coordinate of the electrode, and *σ*= 15 mm is a blurring parameter, selected based on the most conservative estimate of source localization achievable with ECoG (Yekutieli and Benjamini, 1999).

### Electromagnetic field models for TMS

For each stimulation site the TMS coil location and trajectory of the stimulation pulse was recorded with Brainsight across the 50 pulses and averaged, giving a single representative coil position for each anatomical target. This information, along with the intensity of stimulation and coil model was entered into SimNIBS (Thielscher et al., 2015) v 3.0.8. SimNIBS employs finite element modeling with linear basis function to estimate electric field strength in space using individualized models of anatomy derived from each subject’s MRI (Thielscher et al., 2015). This analysis used the headreco model including 6 anatomical layers: white matter volume, gray matter volume, CSF volume, skull volume, skin volume, and eye volumes. The resulting simulated field strength maps were transformed from a SimNIBS-specific volume to each subject’s T1 and MNI152 volumetric space using the same registration techniques described above. This allowed comparison of the E-field strength with the percentage of electrodes at a given distance having a TMS-specific evoked response.

## Results

We start by reviewing safety testing *in vitro* from a phantom brain as well as *in vivo* in humans (N = 20). Next, we discuss human experimental data acquired from 10 neurosurgical participants that received single pulses of TMS delivered to the dlPFC while recording iEEG. Of those subjects, two were eliminated due to excess artifact as described in Methods (*iEEG recording*). Of the remaining eight subjects, we analyzed a total of 1414 electrodes. An overview of the experimental paradigm is shown in Fig. 1a.

**Fig. 1.**
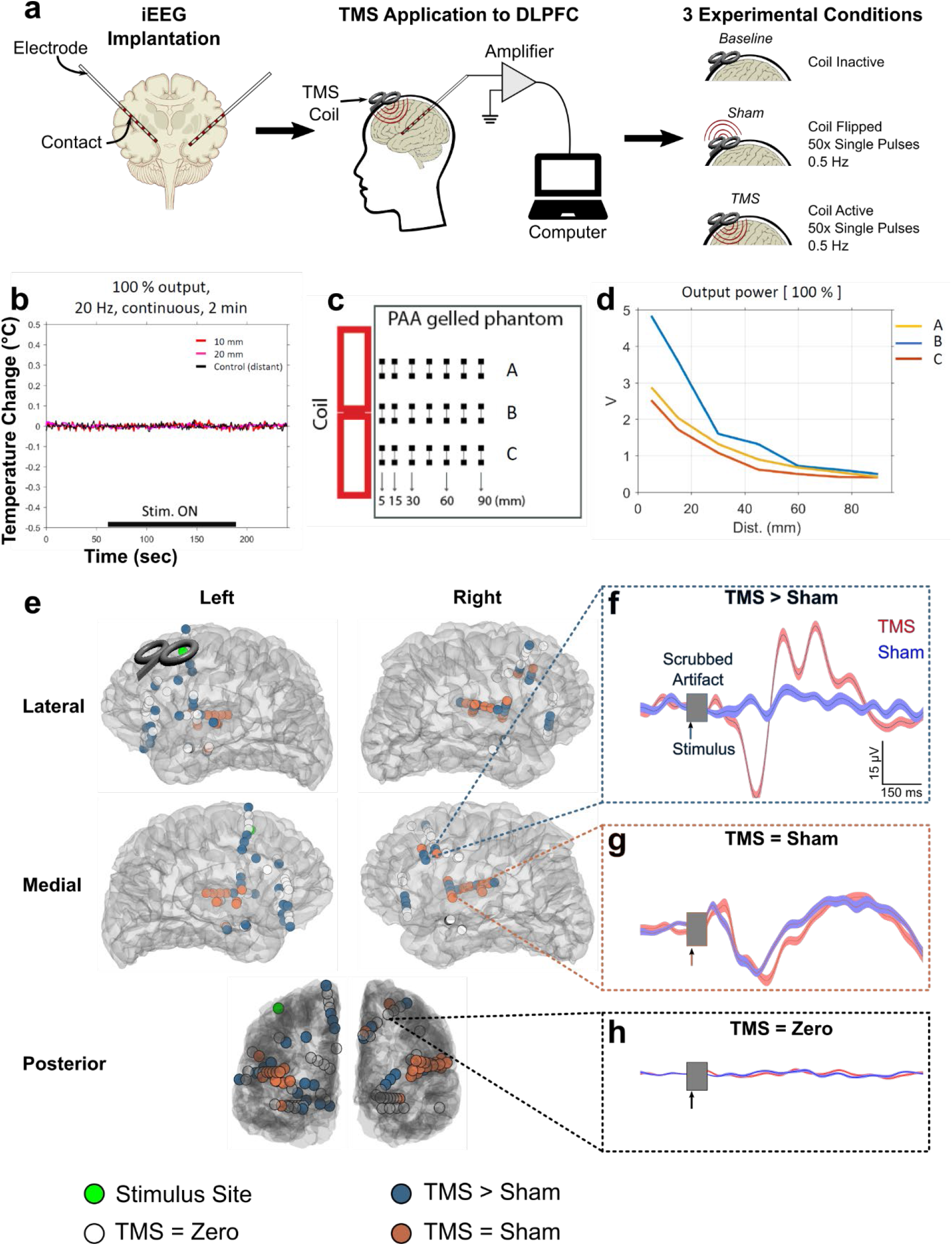
TMS reliably and safely induces intracranial neural responses. **(a)** Schematic of experimental setup. After surgical implantation of iEEG electrodes, subjects received single pulses of TMS while simultaneously recording from iEEG contacts. Two experimental conditions were used: a sham condition with the TMS coil flipped in the opposite direction and a TMS condition with the TMS coil oriented correctly. **(b)** Thermometry traces of temperature of intracranial electrodes while exposed to TMS in an *in vitro* phantom brain. **(c)** Schematic of phantom to study the voltage induced by TMS directed towards intracranial electrodes. Electrodes were placed within a gel phantom in three parallel lines, one at the center of the figure-of-8 coil and the other two each 17.5 mm from the center, aligned along the axis of stimulus delivery. **(d)** Voltage as a function of time to evaluate induced currents. Note that the voltage drops exponentially as a function of distance from the coil both orthogonal and parallel to the axis of stimulation. **(e)** Representative subject’s (Subject 483) brain, with implanted contacts shown as a circle. (f) Representative TMS > sham intracranial TMS evoked potential (iTEP), denoted as *significant* iTEP in the manuscript. For all electrophysiology figures, grey region around time zero represents the time period for which the TMS artifact was removed. Vertical arrow denotes the time when the pulse was delivered. Shaded regions are ±1 SEM. (g) Representative TMS = Sham neural evoked response. (h) Representative electrode without a neural response in either TMS or sham condition.

### TMS is safe in combination with iEEG *in vitro*

First, we evaluated the safety of concurrent TMS-iEEG in a phantom brain model. We observed: 1) no significant heating of electrodes, with all measurements showing minimal change from baseline (<0.1 degree Celsius) (Fig. 1b); 2) no electrode displacement; and 3) the induced voltage within electrodes drops exponentially as a function of distance from the coil both orthogonal and parallel to the axis of stimulation (Fig. 1c,d). Across the various stimulation protocols, we found the maximum voltage induced by TMS was around 5V at 5 mm from the coil when stimulation intensity was set at 100% machine output. This corresponds to a voltage gradient of 0.3 V/mm and a charge density / phase of approximately 7.2 μC/cm^2^, well below the 30 μC/cm^2^ commonly used as a recommended safety threshold for intracranial stimulation (Kuncel and Grill, 2004). Furthermore, these values match the estimated voltage induced by TMS directly within brain tissue (Lu and Ueno, 2017), demonstrating that intracranial electrodes do not cause additional electrical stimulation during TMS.

### Demonstrating safety of TMS-iEEG in humans

20 participants with medically intractable epilepsy enrolled in the study and received TMS while recording concurrently with iEEG. Across all sessions and all stimulation protocols there were no adverse events reported beyond those routinely reported during TMS, such as a worsening of an existing headache or scalp discomfort at the site of stimulation. In those situations when headache or scalp discomfort was reported participants were given options to reduce the stimulation intensity or discontinue the experiment rather than stimulate additional sites. TMS was typically tolerated at 2-4 stimulation sites per participant, with 0.5 Hz being better tolerated than repetitive TMS protocols. There was no change in the frequency of seizures during TMS sessions. A single individual with hundreds of seizures per day each lasting a few seconds had four seizures during one TMS session, which was not different than baseline seizure frequency during the hospitalization. For this patient two seizures occurred during set-up, one during sham stimulation, and one during 0.5 Hz stimulation, at which point the session was discontinued.

### TMS-evoked potentials observed with TMS-iEEG were specific to stimulation

Ten participants received single pulses of TMS delivered to the dlPFC at 0.5Hz. Of these, two participants were excluded from analysis due to significant TMS-related amplifier saturation observed in >10% of contacts (see Methods for details). A typical subject’s intracranial response to TMS is depicted in Fig. 1e with data from all other subjects shown in Supplementary Fig. 1. As can be seen, we were able to successfully isolate Intracranial TMS-Evoked Potentials (iTEPs) that were specific to TMS instead of sham (Fig. 1f). This TMS>sham analysis allowed us to isolate iTEPs while controlling for the auditory responses. Contacts generally fell into three categories: 1) responsive (eliciting a strong iTEP) to TMS specifically over sham (TMS > sham; 8.7% of contacts; Fig. 1f), 2) responsive to both TMS and sham (TMS = sham; 5.8% of contacts; Fig. 1g), and 3) not responsive to either condition (TMS = baseline; 85.3% of contacts; Fig. 1h). Notably, contacts responding to both TMS and sham were enriched in auditory regions such as the left and right transverse temporal cortex (100% and 88%, respectively), suggesting that the TMS = sham condition was effective in controlling for the auditory evoked responses associated with TMS delivery.

### TMS to the dlPFC induces local brain responses in a pattern predicted by the simulated electric field

To evaluate local effects of TMS at a group level we generated a unified coordinate system centered on the stimulation coil, such that electrode locations relative to the coil were combined across subjects. Fig. 2 depicts these results, where all coregistered contacts within 30 mm of the stimulus site (total of 37 contacts) are plotted from the eight analyzed subjects. We first found that in general, significant iTEPs (TMS > sham) were observed in only 18% of contacts within the dlPFC (Fig. 2a). Next, we show that regions across the entire brain with electrodes exposed to a higher electric field were statistically more likely to exhibit significant iTEPs (TMS > sham) (Fig. 2a-d, r = 0.44, p <0.001). In summary, we demonstrate that only a small subset (18%) of electrodes near the dlPFC exhibit significant iTEPs, which can partially be explained by the positive relationship between iTEPs and electric field strength.

**Fig. 2:**
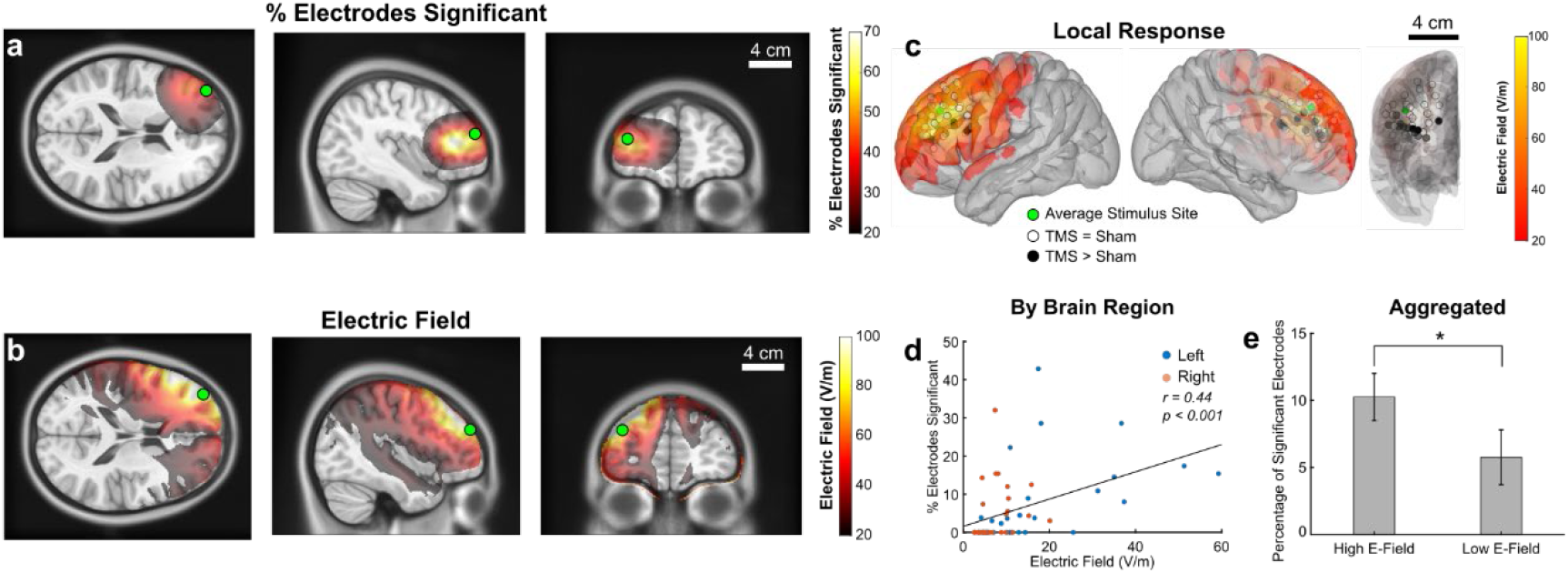
TMS induces local evoked potentials within the dlPFC that correlate with electrical field strength. **(a)** Group plot of realigned electrodes around the average TMS site. Black and white electrodes denote significant (TMS>sham) and non-significant (TMS=sham) intracranial TMS-evoked potentials (iTEPs), respectively. The simulated electric field strength is displayed as a heatmap projected onto the cortical surface. (b) Relationship between average electric field within an anatomic region (as defined in the DKT-Atlas) and the percentage of electrodes with a neural response during TMS. Each dot represents one DKT-defined brain region, color-coded by hemisphere. (c) Merged image of significant iTEPs and electric field strength. (D-E) Relationship between iTEPs and electric field strength. (d)Aggregated average percentage of electrodes in regions above and below the median induced electric field across the brain (High and Low E-Field respectively). *:< 0.05 by Student’s 2-Sample T-Test. (e) Scatter plot of mean iTEPs and electric field strength across brain regions.

### TMS induces network level brain responses in the ACC and insular cortex

TMS is presumed to be therapeutic through network level effects that preferentially modulate sites connected to the stimulation site. To evaluate this possibility, we visualized regional patterns of significant iTEPs (TMS>sham) at the group level based on the proportion of significant iTEP electrode contacts in regions with at least 10 contacts (see Methods; Fig. 3a). In general, there was sufficient coverage across cortical structures, with the exception of the occipital lobe (Supplementary Fig. 2). Ipsilateral to TMS, regions with the highest proportion of significant iTEPs were the dorsal ACC (dACC, 30%), pars opercularis (25%), insula (20%), and middle frontal gyrus (15%). Contralateral to TMS, dACC demonstrated the most consistent iTEPs (30%) with other regions (pars opercularis, temporal and frontal gyri) exhibiting significant iTEPs in <15% of electrodes measured. Regions with a high proportion of significant iTEPs are visualized both on the brain’s transparent surface (Fig. 3c) and MRI slices (Fig. 3e). In these views, iTEPs can be observed in high proportion in the dACC extending superiorly into the dorsomedial prefrontal cortex. In summary, we determined that regional patterns of significant iTEPs following dlPFC TMS were observed in bilateral dACC and ipsilateral insula, pars opercularis, and frontal gyri.

**Fig. 3:**
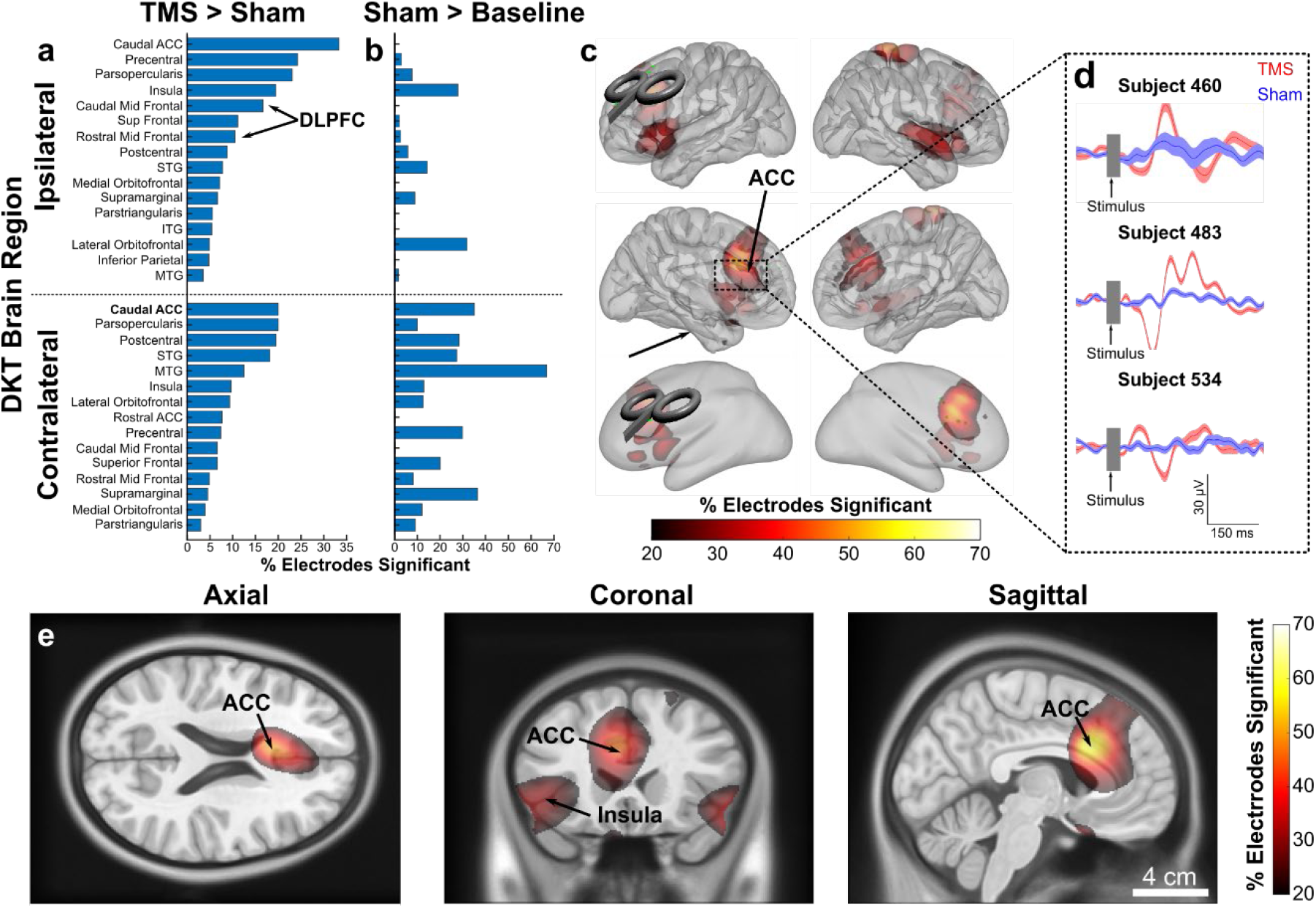
TMS reliably evokes downstream iTEPs within the ACC and Insula. **(a & b).** Bar chart of percentage of electrodes within a Cortical Parcellation (see Methods) that showed a significant iTEP (left; TMS > Sham) as well as response to sham TMS (right; sham>baseline, indicating regions likely showing an auditory response). Regions are split based on ipsilateral and contralateral to the TMS site (top and bottom, respectively). **(c)** Heat maps depicting the percentage of local electrodes that expressed a significant iTEP (TMS > Sham). TMS coil and underlying green dot denote the TMS stimulus site. Note the consistent neural responses in ACC. **(d)** Example iTEPs within the ACC evoked after TMS of the dlPFC. (e) Heat map overlayed depicting the percentage of ACC electrodes that expressed a significant iTEP specifically during TMS.

### Network level iTEPs relate to functional connectivity of the stimulation site

To evaluate whether the pattern of significant evoked responses was related to the functional connectivity of the stimulation site we performed a resting state functional connectivity MRI (rs-fcMRI) analysis seeded from the stimulation site. This was performed both with resting state functional MRI data from the individual participants as well as from a large normative cohort (Fig. 4a; N=98; see Methods), which both had similar patterns of dlPFC-seeded rs-fcMRI connectivity (spatial correlation, Pearson’s r = 0.74, p <0.001). When focusing on regions with the highest percentage of significant iTEPs such as the dACC / dorsomedial prefrontal cortex, we observed robust rs-fcMRI connectivity between dlPFC stimulation site and these regions (Fig. 4a, outlined in black). Across brain regions, those with significant iTEPs demonstrated significantly higher rs-fcMRI connectivity with the dlPFC stimulation site compared to those without significant iTEPs (T_(+)iTEPs_ = 2.6; T_(-)iTEPs_ = 0.8; p <0.001; Fig. 4b). In summary, downstream regions exhibiting dlPFC-induced iTEPs generally exhibited stronger functional connectivity with the dlPFC relative to regions without significant iTEPs.

**Fig. 4:**
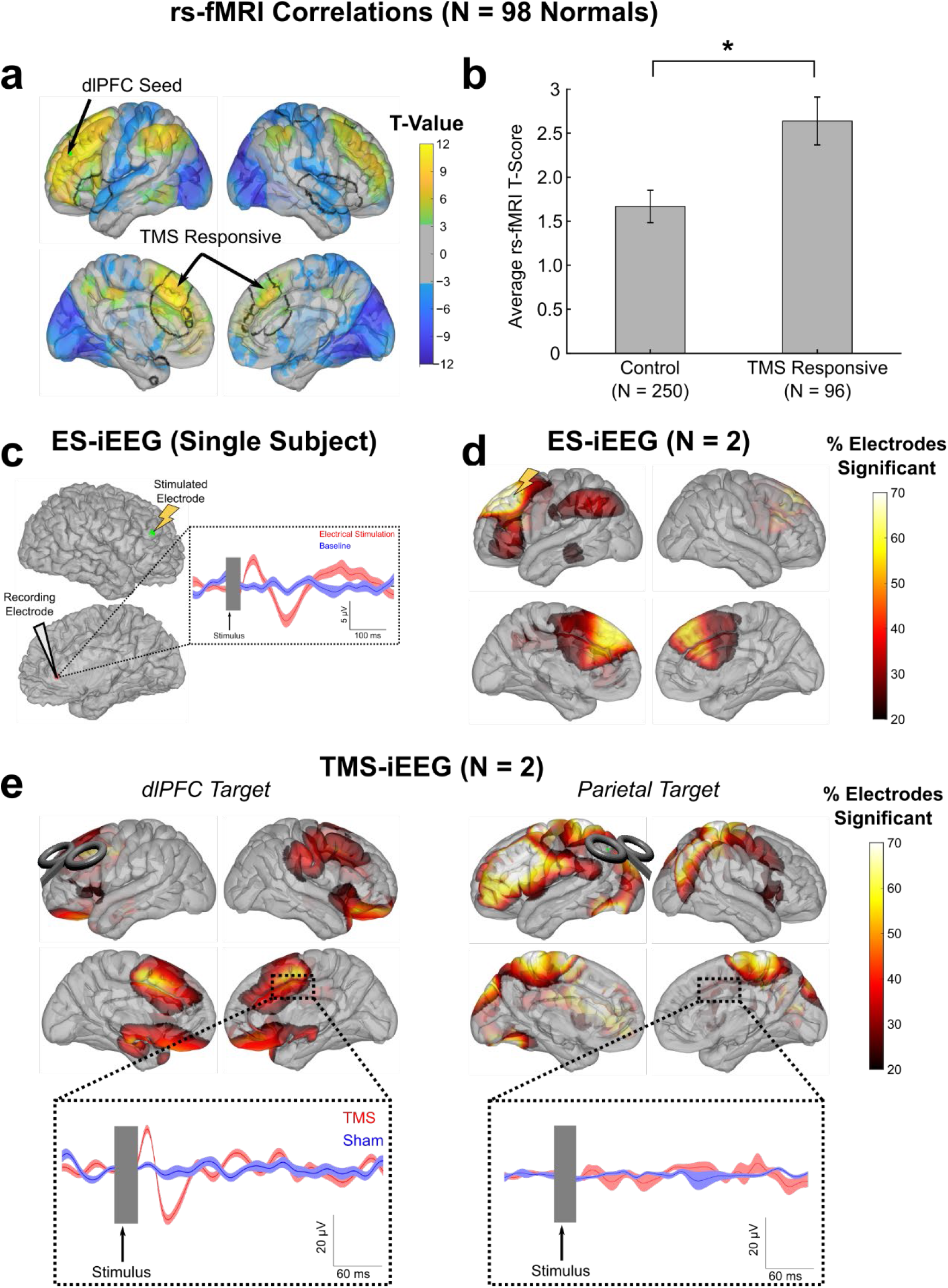
dlPFC TMS activates the ACC in a functionally connected and site-specific manner. **(a)** Resting state functional MRI (rs-fcMRI) maps (N = 98 healthy controls), with a seed determined by the average TMS induced electric field. Depicted are T-Values with FSL’s implementation of nonparametric clustering for multiple comparison correction, with a Z-Stat cutoff of 3.1 (p < 0.001). **(b)** Comparison of rs-fcMRI connectivity values in regions with and without iTEPs. * indicates statistical significance (t = 2.66(344); p=0.0083, 2-sample Student’s T-Test). Error bars are ±1 SEM. **(c-d)** Direct electrical stimulation results. **(c)** Example of iEEG response in the cingulate cortex following single pulses of electrical stimulation to the dlPFC. **(d)** Distribution of evoked potentials after electrical stimulation of the dlPFC (N=2; electrode MNI coordinates: Subject 561: -41, 6, 55; Subject 593: -33, 31, 48). Percent of significant electrodes are plotted on the brain’s surface. Note the significant responses locally and in bilateral medial prefrontal cortex, including ACC. **(e)** Heat map of electrodes that expressed an iTEP specifically after TMS of the dlPFC (*Left*) and the parietal region (*Right*) in the same two subjects. Insert region is the ACC and insert time traces (*bottom*) are from the same electrode. Green dot represents the parietal stimulation site.

### Evaluating potential confounds

One concern is that the ACC iTEP response profile observed in this study is not specific to dlPFC TMS but rather a general response that could be observed with TMS applied to any location. To evaluate this possibility, we performed several additional analyses. First, as pain activates the ACC (Dowdle et al., 2019), we considered the possibility that pain from the TMS discharge and not direct stimulation effects could induce the evoked response pattern observed in the ACC. As direct *electrical* stimulation is not reported to be painful (and often not even perceived), observation of evoked potentials in the ACC after electrical stimulation to the dlPFC would strengthen the argument that iTEPs are produced from cortical propagation from the dlPFC stimulation site. Thus, we applied direct electrical stimulation to the dlPFC and measured ACC evoked responses in a subset of patients with electrode coverage at both locations (Fig. 4c,d; N=2, see Methods). Following single pulse electrical stimulation to the dlPFC, we observed significant electrically-evoked potentials in the medial prefrontal cortex (including ACC regions) as well as parietal lobe. This analysis supports the notion of TMS-induced ACC response due to the propagated effects of stimulation rather than non-specific pain or somatosensory effects.

To evaluate the site specificity of dlPFC TMS on the iTEP response profile in the ACC, in a subset of patients (N=2) we applied single pulses of TMS (0.5Hz) to the inferior parietal lobe (see Methods). Although parietal lobe stimulation induced strong iTEPs in the lateral prefrontal cortex – demonstrating remote iTEPs after parietal TMS – ACC iTEP response profile was not observed (Fig. 4e). It is worth noting that the ACC iTEP response profile was observed in these same patients following dlPFC TMS (Fig. 4e). These results support the notion of anatomical specificity of remote effects of dlPFC TMS. Together, these analyses support the notion that ACC responses following dlPFC TMS is specific to direct stimulation of the dlPFC and not due to pain or somatosensory perceptual changes.

## Discussion

### Summary of findings

After demonstrating safety using a phantom brain model, we performed TMS-iEEG in 20 participants without any observed adverse events. Further, we showed that the proportion of significant local responses from a pulse of TMS relates to the strength of the simulated electromagnetic field. We show that TMS induces brain responses at distant brain sites that were functionally connected to the stimulation site, including the dACC and adjacent medial prefrontal cortex. We further demonstrated that these remote brain responses were specific to dlPFC TMS and not a control site in the parietal lobe, and that dACC responses after direct electrical stimulation of the dlPFC support the notion that these remote responses are unlikely related to the auditory, somatosensory, or pain response to TMS. Taken together, these findings suggest that TMS recruits neuronal populations locally and downstream in functionally connected regions. Work presented here can be taken as evidence for the safety and promise of TMS-iEEG as a new method for interrogating the mechanisms of TMS in humans with high spatiotemporal precision.

### Network-level modulation of the ACC

Prior to this work, data existed to suggest that repetitive TMS to the dlPFC modifies both the dlPFC and a network of connected regions including the ACC and adjacent medial prefrontal structures. Evidence of this has been derived from EEG (Hadas et al., 2019; Kito et al., 2017; Ridder et al., 2011), structural MRI (Boes et al., 2018c; Lan et al., 2016), and fMRI (Baeken et al., 2014; Fox et al., 2012; Ge et al., 2020; Tik et al., 2017). However, without a direct link to intracranial neurophysiology, it has been difficult to confirm the nature of these remote ACC responses. Specifically, it is difficult to source localize subregions of the ACC using EEG and resolve millisecond temporal relationships in the ACC using fMRI. In our study, the dACC was the node with the highest proportion of significant neural response following TMS compared to sham pulses. The dACC also demonstrated strong resting state functional connectivity to the dlPFC and exhibited evoked responses to direct electrical stimulation of the dlPFC, together suggesting a strong and causal connection between the dlPFC and dACC.

A critical question in the field is if therapeutic TMS for depression elicits its clinical effects locally at the dlPFC or downstream in regions functionally connected to the dlPFC, such as the ACC, and if so, which functional subunit of the ACC (dorsal vs subgenual) elicits the clinical effect. Our results demonstrating a strong and causal dlPFC-dACC functional connection supports the enticing notion that modulation of the dACC may play a role in clinical improvement after TMS. Indeed, the dACC is a critical node in the salience network, which controls valence-driven behavior, is activated by negative emotions (Etkin et al., 2011), and exhibits decreased gray matter (Goodkind et al., 2015), and decreased metabolism (Bench et al., 1992; Drevets et al., 1997; George et al., 1997; Ito et al., 1996; Mayberg, 1997) in depression compared to healthy controls. Moreover, electrical stimulation in this region can evoke positive emotion (Bijanki et al., 2019). The more ventral subgenual ACC (sgACC) also has strong evidence relating its activity to depression. In contrast to the dACC, the sgACC tends to have increased metabolism in depression (Ebert et al., 1991; Mayberg, 1997; Wu et al., 1999, 1992), which normalizes after treatment (Mayberg, 1997), and appears to play a role in ruminations characteristic of depression (Berman et al., 2011; Grimm et al., 2009; Sheline et al., 2009). The pattern of resting fMRI connectivity between the dlPFC and these two ACC regions is different, with dlPFC activity positively correlated with dACC activity (Chen et al., 2013) and negatively correlated with sgACC activity (Fox et al., 2012). A major question in the field is if TMS to the dlPFC modulates both the dACC and sgACC directly, or if the effects are direct at one site and indirect at another. While our electrode coverage was greater at the dACC relative to the sgACC (see Supplementary Fig. 2), our results to date support a causal dlPFC-dACC connection elucidated by single pulses of TMS. Whether a similar propagation pattern exists for sgACC in response to dlPFC TMS will require further study with denser sampling of the sgACC.

### Limitations and future directions

This analysis has limitations, some of which can be addressed in future experiments. First, the TMS artifact saturated the iEEG amplifiers and degraded the physiological signal within the first 15 milliseconds after TMS. To account for this, we focused our analyses to start 15 milliseconds after single pulses of TMS were administered. While this strategy minimized the potential for TMS artifact contaminating the physiological signal, it limited our analysis of the immediate (<15 ms) effects of TMS. This may explain why iTEPs were only observed in 18% of electrodes immediately local to the stimulus site. This may be partially overcome in future analyses through enhanced artifact rejection strategies and through new amplifiers that accommodate a wider input range such that TMS does not saturate the signal. Second, our sample size was small and patients were heterogeneous with respect to seizure onset, electrode type, and the distribution of anatomical coverage. Therefore, findings from this study may be skewed towards regions with greater anatomical coverage and miss regions with robust responses but without coverage. A larger study will be necessary to further explore iTEPs in these undersampled regions. Finally, these experiments were conducted on patients with medication refractory epilepsy taking anti-epileptic medication. Although electrodes in the epileptic network were removed (see Methods), the seizure focus and early epileptic spread regions and seizure medications can impact local and global brain excitability and connectivity (Bettus et al., 2011; Pereira et al., 2010; Pittau et al., 2012). Thus, further study is needed to determine how these responses may differ from healthy participants not on medications.

## Conclusions

Taken together, these results provide compelling proof-of-concept results to suggest that the physiological effects of TMS can be recorded with intracranial electrodes in humans. We observed no adverse effects of TMS-iEEG experiments in twenty participants to date. While encouraging, extreme caution must be taken to ensure continued patient safety. We are optimistic TMS-iEEG will provide an informative novel methodology in the ongoing efforts to understand the underlying mechanisms of TMS.

## Acknowledgements

First, we thank the neurosurgery patients who volunteered in this research. We also thank Christopher Kovach, Ariane Rhone, Haiming Chen, and Benjamin Pace for their assistance with image processing, coordinating, and conducting the experiments. This research was supported by NIMH R21MH120441 and 5R01dC004290-20. J.B.W. was supported by F30MH119763 and the Mark and Mary Stevens Interdisciplinary Graduate Fellowship. C.J.K was supported by R01MH126639, R01MH129018, and a Burroughs Wellcome Fund Career Award for Medical Scientists. A.D.B. was also supported by R01NS114405. N.T.T. was supported by 5T32-MH019113. This work was conducted, in part, on an MRI instrument funded by 1S10OD025025-01.

## Author contributions

JBW, JEB, HO, CJK, and ADB conceived of the study. JBW performed electrophysiology analyses. JEB performed imaging and electric field simulation analyses. HO performed safety testing and analyses and participated in experimental testing with TMS and intracranial stimulation. BDU, NTT, and PEG participated in experimental testing. MH helped with study design, safety, and subject recruitment. All authors contributed to writing and editing the manuscript. ADB and CJK jointly supervised all aspects of the study. All authors have seen and approved the manuscript, and it has not been accepted or published elsewhere.

## Competing interests

The authors declare no competing interests. CJK currently holds equity in Alto Neurosciences, Inc.

**Supplementary information is available separately.**

**Correspondence and requests for materials** should be addressed to Aaron Boes or Corey Keller.

