## Supplemental Materials for "Effects of transcranial magnetic stimulation on the human brain recorded with intracranial electrocorticography: First-in-human study"

**Supplementary Figures and Tables**

| ID | Age  (yr) | Sex | Handed-ness | Ethnicity | Education (yr) | Age of Onset (yr) | Onset of Epilepsy | Relevant Co-Morbidities | Lesions | Prior Surgical Resection | Motor Threshold | Day of Testing Medications |
| --- | --- | --- | --- | --- | --- | --- | --- | --- | --- | --- | --- | --- |
| 403 | 56 | F | R | non-Hispanic White | 15 | 19 | left anterior temporal | hypothyroidism; depression; anxiety | none | No | 70% | biotin, cholecalciferol, clonazepam, diazepam, fluoxetine HCl, hydroxyzine HCl, lamotrigine, levothyroxine sodium, magnesium oxide, metoclopramide HCL, oxcarbazepine, perampanel |
| 404 | 44 | F | R | non-Hispanic White | 13 | 14 | left parietal | leukocytosis | none | No | 49% | clonazepam, lacosamide, levetiracetam |
| 405 | 19 | M | R | non-Hispanic White | 12 | 9 | left frontal | left frontal hemorrhage (age 9) | left frontal | No | 88% | acetaminophen, bisacodyl, cefazolin, docusate, heparin, levetiracetam, Lorazepam, magnesium hydroxide, morphine, omeprazole, ondansetron, oxcarbazepine, perampanel, sennosides, sodium chloride 0.9% |
| 409 | 31 | F | R | non-Hispanic White | 11 (GED) | 25 | left anterior temporal | bipolar I | none | No | 78% | acetaminophen, caffeine, bisacodyl, clonazepam, ferrous sulfate, hydromorphone, hydroxyzine pamoate, lacosamide, levetiracetam, lurasidone, melatonin, omeprazole |
| 416 | 34 | M | A | non-Hispanic White | 12 | 20 | left occipital | TBI motorcycle accident | focal cortical dysplasia type IIb | No | 59% | lacosamide, levetiracetam, cannabidiol oil |
| 423 | 51 | M | R | non-Hispanic White | 8 | * | left anterior temporal | VNS stimulator | none | No | 49% | albuterol sulfate, carbamazepine, fluticasone, propionate, guaifenesin/dextromethorphan, ibuprofen, mirtazapine, tiotropium bromide, zonisamide |
| 429 | 33 | F | R | non-Hispanic White | 14 | 20 | right anterior temporal | depression | none | No | 86% | albuterol sulfate, folic acid, lorazepam, norgestimate-ethinyl estradiol, sertraline HCl |
| 430 | 28 | M | R | non-Hispanic White | 12 | 4 | generalized | none | none | No | 53% | clonazepam, lamotrigine, risperidone, topiramate, venlafaxine HCl |
| 460 | 52 | M | R | non-Hispanic White | 12 | 16 | left medial temporal | anxiety, depression | mild cortical atrophy and chronic small vessel disease | No | 48% | aspirin, cetirizine HCl, doxazosin mesylate, escitalopram oxalate, lamotrigine, losartan potassium, magnesium chloride, riboflavin, sennosides, tamsulosin HCl |
| 477 | 23 | F | R | non-Hispanic White | 12 | 16 | left lateral posterior parietal | moderate persistent asthma | none | No | 59% | albuterol sulfate, clonazepam, fluticasone propion/salmeterol, ipratropium/albuterol sulfate, levetiracetam, montelukast sodium, oxcarbazepine, sertraline HCl, topiramate |
| 483* | 18 | M | R * | non-Hispanic White | 12 | 3 | generalized | Type I Diabetes | none | No | 75% | cefazolin, clobazam, felbamate, Insulin, miralax |
| 493 | 35 | M | L | non-Hispanic White | 16 | 7 | right medial temporal & orbitofrontal | none | none | No | 69% | levetiracetam, lamotrigine and oxcarbazepine |
| 518 | 14 | F | R | non-Hispanic White | 11 | 8 | right posterior-temporal | none | right occipital, temporal, and parietal resections | Yes | n/a | clonazepam, lamotrigine, topomax |
| 524 | 18 | M | R | non-Hispanic White | 12 | 16 | right frontal, bilateral medial temporal | suicidality, hypersomnia | none | No | 88% | oxycodone, zonisamide, dextroamphetamine, escitalopram, midazolam, lithium, modafinil, nortriptyline, propanolol |
| 534 | 20 | M | R | non-Hispanic White | 12 | 11 | generalized | none | amygdalohipp-ocampectomy one year prior | Yes | n/a | clobazam, oxcarbazepine |
| 538 | 19 | F | R | non-Hispanic White | 12 | 11 | left medial temporal | mild depression | left amygdalohipp-ocampectomy three years prior | Yes | 80% | buspirone, cetirizine, diphenhydramine, docusate, famotidine, levetiracetam, lorazapam, vancomyzinec |
| 559 | 27 | F | R | non-White | 12 | 18 | right fronto-  temporal | depression; anxiety | frontal cortical dysplasia | No | n/a | carbamazepine, clonazepam, lacosomide, levetiracetam, ondansetron, sumatriptan, zonisamide |
| 561 | 19 | M | R | non-Hispanic White | 12 | 9 | right centroparieto-occipital | allergic rhinitis, primary insomnia, adolescent scoliosis, MDD | none | No | n/a | acetaminophen, carbamazepine, cefazolin, cetirizine, docusate, ibuprofen, lacosamide, lorazepam, ondansetron, oxycodone, miralax, sertraline |
| 579 | 13 | F | * | non-Hispanic White | 8 | childhood | bilateral generalized | ocular cyst (removed) | periventricular gray matter heterotopias | No | n/a | clobazam, clonazepam, fluoxetine, lacosimide, midazolam, oxcarbazepine |
| 593 | 15 | F | R | non-Hispanic White | 9 | 1 | left temporal | mild malnutrition; left-sided intraparenchymal hemorrhage | left-sided intraparenchy-mal hemorrhage | No | 55% | lacosamide, lamotrigine, valproic acid, levetiracetam, cefazolin |

**Supplementary Table 1. Patient Demographics, Epilepsy Etiology, Lesions, Co-morbidities, Motor threshold, and Medications**

| DKT Label | # Contacts | # Patients | | % Sig. | % Patients | Avg. T-Value | Avg. P-Value |
| --- | --- | --- | --- | --- | --- | --- | --- |
| Ctx-lh-bankssts | 8 | 2 | 0.00 | | 0.00 | 39.85 | 0.54 |
| Ctx-lh-caudalanteriorcingulate | 10 | 3 | 0.33 | | 1.00 | 338.13 | 0.01 |
| Ctx-lh-caudalmiddlefrontal | 18 | 4 | 0.17 | | 0.25 | 126.15 | 0.40 |
| Ctx-lh-cuneus | 1 | 1 | 0.00 | | 0.00 | 69.05 | 0.08 |
| Ctx-lh-entorhinal | 4 | 2 | 0.00 | | 0.00 | 240.46 | 0.01 |
| Ctx-lh-fusiform | 23 | 6 | 0.00 | | 0.00 | 41.94 | 0.57 |
| Ctx-lh-inferiorparietal | 42 | 4 | 0.05 | | 0.25 | 52.93 | 0.59 |
| Ctx-lh-inferiortemporal | 37 | 5 | 0.05 | | 0.40 | 110.54 | 0.49 |
| Ctx-lh-insula | 36 | 5 | 0.19 | | 0.60 | 274.00 | 0.22 |
| Ctx-lh-isthmuscingulate | 15 | 4 | 0.00 | | 0.00 | 58.07 | 0.40 |
| Ctx-lh-lateraloccipital | 14 | 2 | 0.00 | | 0.00 | 29.05 | 0.75 |
| Ctx-lh-lateralorbitofrontal | 41 | 5 | 0.05 | | 0.40 | 119.07 | 0.26 |
| Ctx-lh-lingual | 16 | 4 | 0.00 | | 0.00 | 43.24 | 0.59 |
| Ctx-lh-medialorbitofrontal | 14 | 4 | 0.07 | | 0.25 | 126.22 | 0.43 |
| Ctx-lh-middletemporal | 57 | 6 | 0.04 | | 0.33 | 41.79 | 0.63 |
| Ctx-lh-paracentral | 6 | 3 | 0.00 | | 0.00 | 503.18 | 0.22 |
| Ctx-lh-parahippocampal | 8 | 4 | 0.00 | | 0.00 | 12.61 | 0.63 |
| Ctx-lh-parsopercularis | 13 | 4 | 0.23 | | 0.25 | 159.20 | 0.38 |
| Ctx-lh-parsorbitalis | 6 | 3 | 0.33 | | 0.67 | 102.05 | 0.51 |
| Ctx-lh-parstriangularis | 18 | 4 | 0.06 | | 0.25 | 77.54 | 0.34 |
| Ctx-lh-pericalcarine | 3 | 1 | 0.00 | | 0.00 | 22.23 | 0.49 |
| Ctx-lh-postcentral | 34 | 5 | 0.09 | | 0.40 | 199.76 | 0.26 |
| Ctx-lh-posteriorcingulate | 7 | 3 | 0.00 | | 0.00 | 65.62 | 0.45 |
| Ctx-lh-precentral | 33 | 5 | 0.24 | | 0.60 | 348.75 | 0.14 |
| Ctx-lh-precuneus | 30 | 5 | 0.00 | | 0.00 | 68.22 | 0.30 |
| Ctx-lh-rostralanteriorcingulate | 11 | 3 | 0.00 | | 0.00 | 181.32 | 0.11 |
| Ctx-lh-rostralmiddlefrontal | 38 | 4 | 0.11 | | 0.25 | 98.75 | 0.45 |
| Ctx-lh-superiorfrontal | 45 | 4 | 0.11 | | 0.50 | 132.39 | 0.36 |
| Ctx-lh-superiorparietal | 40 | 5 | 0.00 | | 0.00 | 68.97 | 0.43 |
| Ctx-lh-superiortemporal | 77 | 6 | 0.08 | | 0.33 | 134.25 | 0.55 |
| Ctx-lh-supramarginal | 45 | 5 | 0.07 | | 0.40 | 110.10 | 0.35 |
| Ctx-lh-temporalpole | 5 | 1 | 0.00 | | 0.00 | 66.98 | 0.61 |
| Ctx-lh-transversetemporal | 8 | 3 | 0.25 | | 0.67 | 238.00 | 0.02 |
| Ctx-lh-unknown | 4 | 4 | 0.00 | | 0.00 | 73.66 | 0.53 |
| Ctx-rh-caudalanteriorcingulate | 20 | 3 | 0.20 | | 0.33 | 163.32 | 0.35 |
| Ctx-rh-caudalmiddlefrontal | 15 | 3 | 0.07 | | 0.33 | 86.11 | 0.60 |
| Ctx-rh-cuneus | 1 | 1 | 0.00 | | 0.00 | 0.00 | 1.00 |
| Ctx-rh-entorhinal | 6 | 1 | 0.50 | | 1.00 | 218.38 | 0.01 |
| Ctx-rh-fusiform | 11 | 3 | 0.00 | | 0.00 | 2362.44 | 0.55 |
| Ctx-rh-inferiorparietal | 15 | 2 | 0.00 | | 0.00 | 62.14 | 0.46 |
| Ctx-rh-inferiortemporal | 11 | 4 | 0.00 | | 0.00 | 49.61 | 0.44 |
| Ctx-rh-insula | 31 | 5 | 0.10 | | 0.20 | 288.09 | 0.12 |
| Ctx-rh-isthmuscingulate | 2 | 1 | 0.00 | | 0.00 | 88.01 | 0.08 |
| Ctx-rh-lateraloccipital | 6 | 2 | 0.00 | | 0.00 | 22.67 | 0.58 |
| Ctx-rh-lateralorbitofrontal | 64 | 6 | 0.09 | | 0.33 | 141.10 | 0.52 |
| Ctx-rh-lingual | 3 | 1 | 0.00 | | 0.00 | 44.78 | 0.68 |
| Ctx-rh-medialorbitofrontal | 25 | 6 | 0.04 | | 0.17 | 62.02 | 0.45 |
| Ctx-rh-middletemporal | 24 | 4 | 0.13 | | 0.25 | 388.97 | 0.15 |
| Ctx-rh-paracentral | 16 | 2 | 0.00 | | 0.00 | 91.84 | 0.29 |
| Ctx-rh-parsopercularis | 10 | 2 | 0.20 | | 0.50 | 146.69 | 0.28 |
| Ctx-rh-parsorbitalis | 17 | 3 | 0.00 | | 0.00 | 6.28 | 0.90 |
| Ctx-rh-parstriangularis | 33 | 4 | 0.03 | | 0.25 | 44.60 | 0.51 |
| Ctx-rh-postcentral | 46 | 4 | 0.20 | | 1.00 | 174.95 | 0.21 |
| Ctx-rh-posteriorcingulate | 15 | 3 | 0.00 | | 0.00 | 73.55 | 0.30 |
| Ctx-rh-precentral | 94 | 4 | 0.07 | | 0.75 | 191.24 | 0.15 |
| Ctx-rh-precuneus | 10 | 2 | 0.00 | | 0.00 | 54.62 | 0.31 |
| Ctx-rh-rostralanteriorcingulate | 13 | 4 | 0.08 | | 0.25 | 197.07 | 0.34 |
| Ctx-rh-rostralmiddlefrontal | 61 | 3 | 0.05 | | 0.33 | 84.93 | 0.63 |
| Ctx-rh-superiorfrontal | 45 | 3 | 0.07 | | 0.33 | 317.95 | 0.37 |
| Ctx-rh-superiorparietal | 12 | 2 | 0.00 | | 0.00 | 69.51 | 0.46 |
| Ctx-rh-superiortemporal | 11 | 3 | 0.18 | | 0.33 | 192.88 | 0.13 |
| Ctx-rh-supramarginal | 22 | 3 | 0.05 | | 0.33 | 159.31 | 0.32 |
| Ctx-rh-temporalpole | 24 | 1 | 0.00 | | 0.00 | 51.79 | 0.27 |
| Ctx-rh-transversetemporal | 4 | 1 | 0.25 | | 1.00 | 358.17 | 0.00 |
| Ctx-rh-unknown | 2 | 1 | 0.00 | | 0.00 | 0.00 | 1.00 |
| Left-Amygdala | 5 | 2 | 0.00 | | 0.00 | 20.47 | 0.51 |
| Left-Hippocampus | 18 | 5 | 0.00 | | 0.00 | 50.98 | 0.65 |
| Left-VentralDC | 2 | 2 | 0.50 | | 0.50 | 99.78 | 0.51 |
| Right-Amygdala | 16 | 4 | 0.00 | | 0.00 | 122.48 | 0.18 |
| Right-Hippocampus | 23 | 4 | 0.00 | | 0.00 | 121.75 | 0.20 |
| Right-Inf-Lat-Vent | 2 | 2 | 0.00 | | 0.00 | 25.89 | 0.57 |
| Right-Thalamus-Proper | 3 | 1 | 0.00 | | 0.00 | 187.16 | 0.01 |
| Right-VentralDC | 3 | 2 | 0.00 | | 0.00 | 41.07 | 0.18 |
| Unknown | 7 | 5 | 0.00 | | 0.00 | 87.54 | 0.21 |
| Other | 36 | 13 | 0.02 | | 0.03 | 118.73 | 0.44 |

**Supplementary Table 2** **–TMS and sham induced iTEPs at each cortical region.** For each parcellated region, number of patients with at least one electrode in that region and the total number of contacts are shown. Additionally, the percent of contacts, percent of patients, and average T- and P-values from TMS and sham-induced iTEPs, are shown.

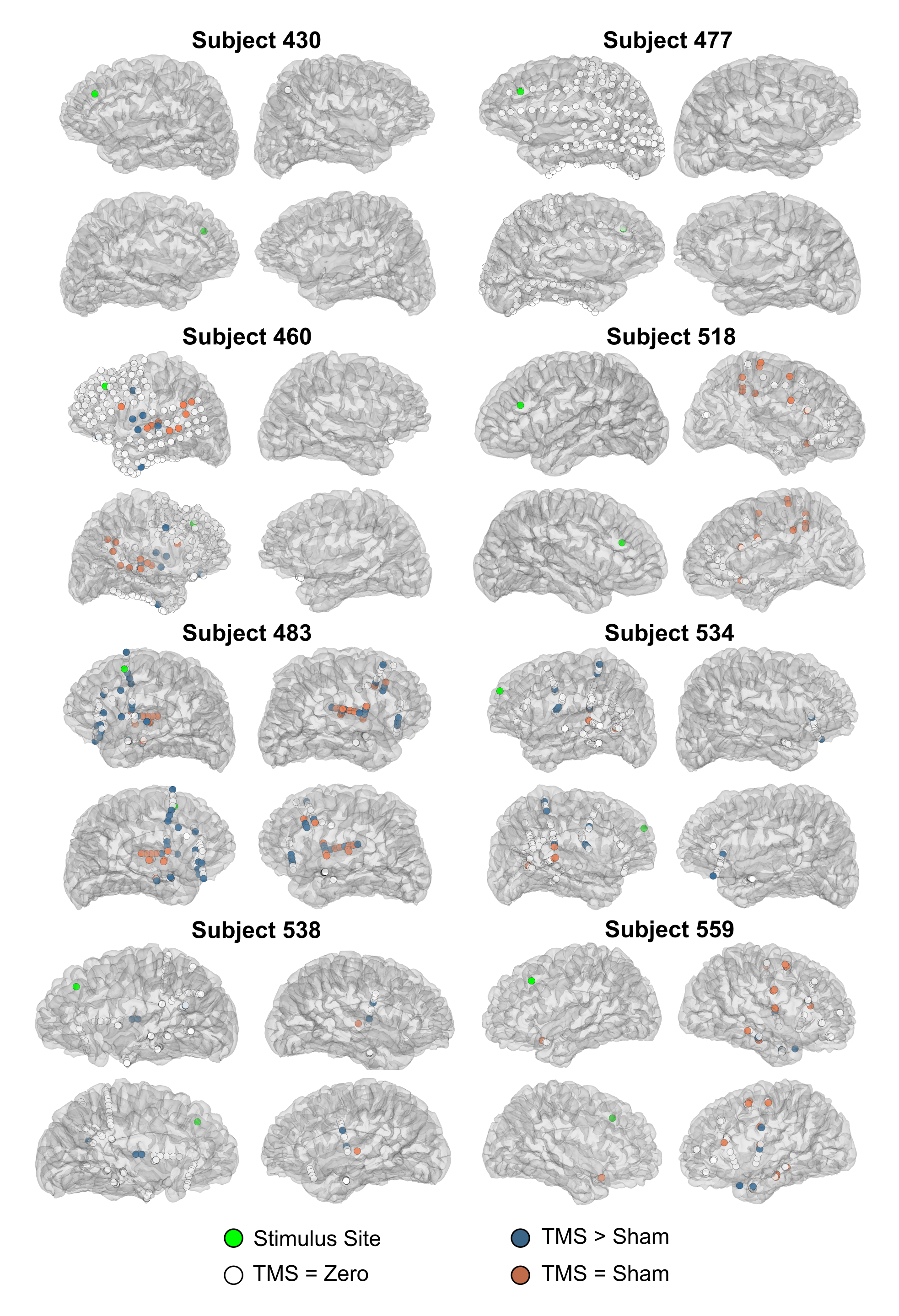

**Supplementary Fig. 1:** Electrode and iTEP coverage for each subject that received dlPFC TMS.

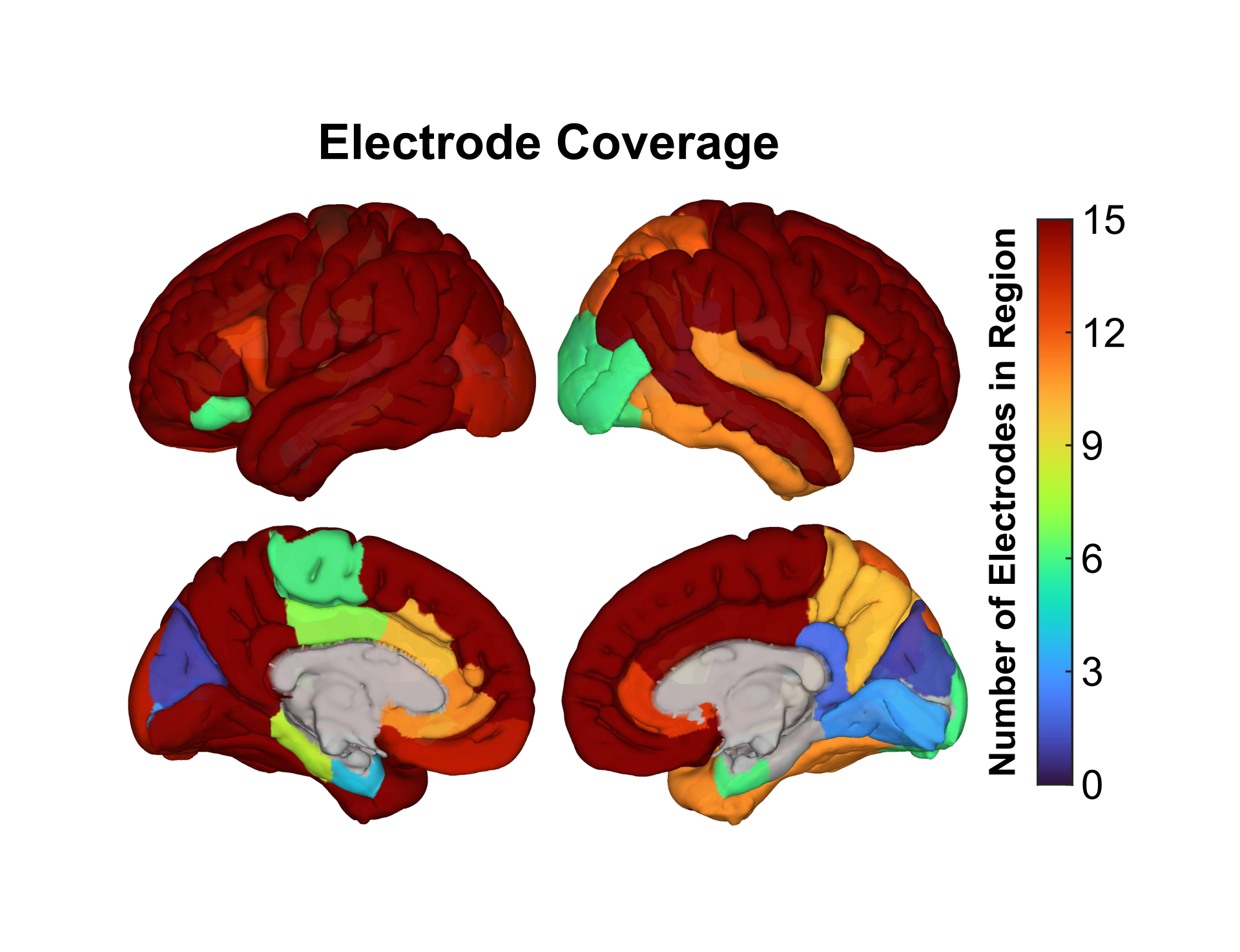

**Supplementary Fig. 2:** Electrode coverage across all 8 subjects, plotted as number of electrodes per brain region, as defined in the DKT atlas.

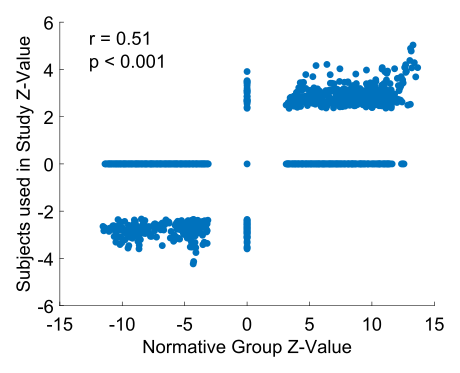

**Supplementary Fig. 3:** Comparison of rs-fcMRI connectivity Z-Values for our normative group (depicted in Fig. 4a; N = 98) and subjects used in our dlPFC analysis (N = 10). Each dot represents one vertex's connectivity to the dlPFC. Correlation is Pearson's Correlation, and p-value is based on 2-tailed Student's T-test.
